# Parcellation of the neonatal cortex using Surface-based Melbourne Children’s Regional Infant Brain atlases (M-CRIB-S)

**DOI:** 10.1101/544304

**Authors:** Chris L. Adamson, Bonnie Alexander, Gareth Ball, Richard Beare, Jeanie L.Y. Cheong, Alicia J. Spittle, Lex W. Doyle, Peter J. Anderson, Marc L. Seal, Deanne K. Thompson

## Abstract

Longitudinal studies measuring changes in cortical morphology over time are best facilitated by parcellation schemes compatible across all life stages. The Melbourne Children’s Regional Infant Brain (M-CRIB) and M-CRIB 2.0 atlases provide voxel-based parcellations of the cerebral cortex compatible with the Desikan-Killiany (DK) and the Desikan-Killiany-Tourville (DKT) cortical labelling schemes. However, there is still a need for a surface-based approach for parcellating neonatal images using these atlases.

We introduce surface-based versions of the M-CRIB and M-CRIB 2.0 atlases, termed M-CRIB-S(DK) and M-CRIB-S(DKT), with a pipeline for automated parcellation utilizing FreeSurfer and developing Human Connectome Project (dHCP) tools.

Using *T*_2_-weighted magnetic resonance images of healthy neonates (*n* = 58), we created average spherical templates of cortical curvature and sulcal depth. Manually-labelled regions in a subset (*n* = 10) were encoded into the spherical template space to construct M-CRIB-S(DK) and M-CRIB-S(DKT) atlases. Labelling accuracy was assessed using Dice overlap measures and boundary discrepancy measures within a leave-one-out cross-validation framework.

Cross-validated labelling accuracy was high for both atlases (average regional Dice = 0.79 - 0.83). Worst-case boundary discrepancy instances ranged from 9.96 - 10.22 mm, which appeared to be driven by variability in anatomy for particular cases.

The M-CRIB-S atlases and automatic pipeline allow extraction of neonatal cortical surfaces labelled according to the DK or DKT parcellation schemes. This will facilitate surface-based investigations of brain structure at the neonatal time point. The atlases and spherical surfaces, along with customised scripts for segmentation, cortical surface extraction and parcellation, are available for public download.

## Introduction

The delineation of cortical areas on magnetic resonance images (MRI) A considered to be a prerequisite for beginning to understand the complexities of the human brain (Brett et al. 2002). Accurate understanding of *development* of the brain is reliant on accurate parcellation of the cerebral cortex, from around the time of normal birth onwards.

*FreeSurfer* (B. Fischl 2012; B. Fischl and Dale 2000; Bruce Fischl et al. 2002; B. Fischl et al. 2004) is a commonly used cortical extraction and parcellation software suite applicable to T1 MRI scans of children and adults, and its available parcellation schemes include the Desikan-Killiany (DK) (Desikan et al. 2006) and Desikan-Killiany-Tourville (DKT) (Klein and Tourville 2012) adult atlases. However, tools tuned for adult brains, such as the adult *T*1-based templates and atlases provided in *FreeSurfer* (B. Fischl et al. 1999; B. Fischl et al. 2004), are not directly applicable to neonatal brain images. Tissue signal intensities are different in neonatal brains compared to those in adults (L. Wang et al. 2019). Thus the optimal MRI sequences for the grey and white matter contrast required to identify cortical surfaces differ by age. While *T*_*1*_-weighting is optimal for adult brains, *T*_*2*_-weighted contrasts are optimal for neonatal brains. Consequently, specialized algorithms are required in order to contend with neonatal-specific tissue intensities (Beare et al. 2016; Makropoulos et al. 2016). Thus, brain atlases and image segmentation and parcellation tools specific for infants are required.

Methods for cortical parcellation of infant brain images have focused on warping standardized cortical atlases from adult brains (Klein and Tourville 2012; Desikan et al. 2006; Tzourio-Mazoyer et al. 2002) onto infant brains e.g. (Shi et al. 2011; Wu et al. 2018). However, labelling a neonatal brain image using adult-derived atlases is problematic, due to the inherent difference in anatomy and tissue composition between infant and adult brains (Richards et al. 2016). Recently, we introduced the Melbourne Children’s Regional Infant Brain (M-CRIB) atlases (Alexander et al. 2017; Alexander et al. 2019b), which are neonatal-specific, voxel-wise brain atlases. The cortical parcellations were constructed to be compliant with the DK (Desikan et al. 2006) and DKT (Klein and Tourville 2012) adult cortical parcellation schemes. Each of the M-CRIB atlases are comprised of 10 neonatal brains that have been extensively manually parcellated to accurately reflect brain structures at this time-point. Parcellation of new data has been achieved using multi-atlas label fusion algorithms that probabilistically assign labels to each voxel after warping the set of parcellated atlases to a novel dataset e.g. (Alexander et al. 2019a; Akhondi-Asl and Warfield 2013).

While accurate labelling can be achieved using voxel-based parcellation schemes (Makropoulos et al. 2018; Alexander et al. 2019a; Alexander et al. 2017), surface-based registration methods lead to significantly improved alignment of cortical landmarks, including cortical folds, and therefore more accurate placement of areal boundaries (Ghosh et al. 2010; Coalson et al. 2018).

The Developing Human Connectome Project (dHCP) has recently provided a pipeline for segmentation and cortical extraction for *T*_2_-weighted images of neonatal brains (Makropoulos et al. 2018). This process segments brain tissue into cerebral and cerebellar grey and white matter, and various subcortical grey matter structures, before extracting an inner and outer cortical surface which is automatically partitioned into lobes. These existing cortical surface extraction tools can be expanded upon to incorporate a neonatal brain atlas and provide accurate surface-based parcellations. Thus, measures such as cortical thickness, surface area, and curvature for each cortical sub-division of the M-CRIB atlases can subsequently be derived for infant MRI scans.

In this study we aimed to provide neonatal average surface templates and surface-based cortical atlases based on the M-CRIB and M-CRIB 2.0 parcellation schemes, that are compatible with *FreeSurfer* and the dHCP pipelines (B. Fischl et al. 2004; B. Fischl 2012; B. Fischl et al. 1999). Additionally, we aimed to provide companion scripts to perform cortical surface extraction, surface registration and atlas-based cortical parcellation using novel neonatal *T*_2_-weighted brain images. Given the compatibility of the neonatal M-CRIB-S parcellation schemes with the adult DK and DKT schemes, the proposed tools can generate parcellated neonatal cortical regions that are comparable with those obtained using existing tools such as FreeSurfer at older time points, enabling longitudinal studies, beginning from the neonatal time-point.

## Methods

### Participants

A total of 58 term-born (≥ 37 weeks’ gestation), healthy neonates (40.2 - 44.9 weeks post-menstrual age (PMA) at scan, *M* = 42.4, *SD* = 1.2, 26 female) were scanned as control subjects as part of preterm birth studies (Spittle et al. 2014; Walsh et al. 2014). Criteria for a subject being healthy were no admissions to a neonatal intensive care or special care unit, resuscitation at birth not required, birthweight more than 2.5 kg and no evidence of congenital conditions known to affect development and growth. Ethical approval for the studies was obtained from the Human Research Ethics Committees of the Royal Women’s Hospital and the Royal Children’s Hospital, Melbourne. Written informed consent was obtained from parents. Data that exhibited excessive movement or other corrupting artefacts were excluded. This cohort was subdivided into the following two subsets: *labelled* and *unlabelled* subsets. The *labelled* set comprised the ten subjects (40.29 – 43.00 weeks’ PMA at scan, *M* = 41.71, *SD* = 1.31, 4 female) of the M-CRIB atlas that had been previously selected from this cohort on the basis of minimal motion or other artifact on the T2 images (Alexander et al. 2019b; Alexander et al. 2017). The *unlabelled* subset consisted of the remaining 48 subjects (40.2 – 44.9 weeks’ PMA at scan, *M* = 42.6, *SD* = 1.3, 22 female).

### MRI acquisition

All neonate subjects were scanned at the Royal Children’s Hospital, Melbourne, Australia, on a 3T Siemens Magnetom Trio scanner during unsedated sleep. *T*_2_-weighted images were acquired with a turbo spin echo sequence with the following parameters: 1 mm axial slices, flip angle = 120°, repetition time = 8910 ms, echo time = 152 ms, field of view = 192 × 192 mm, in-plane resolution = 1 mm^2^ (zero-filled interpolated to 0.5 × 0.5 × 1 mm in image reconstruction), matrix size = 384 × 384. All *T*_2_-weighted images were resliced to voxel-volume-preserving size of 0.63 × 0.63 × 0.63 mm (Loh et al. 2016; Alexander et al. 2017).

### Processing pipeline

The proposed processing pipeline and M-CRIB-S training data is graphically described in Figure 1.

**Figure 1:**
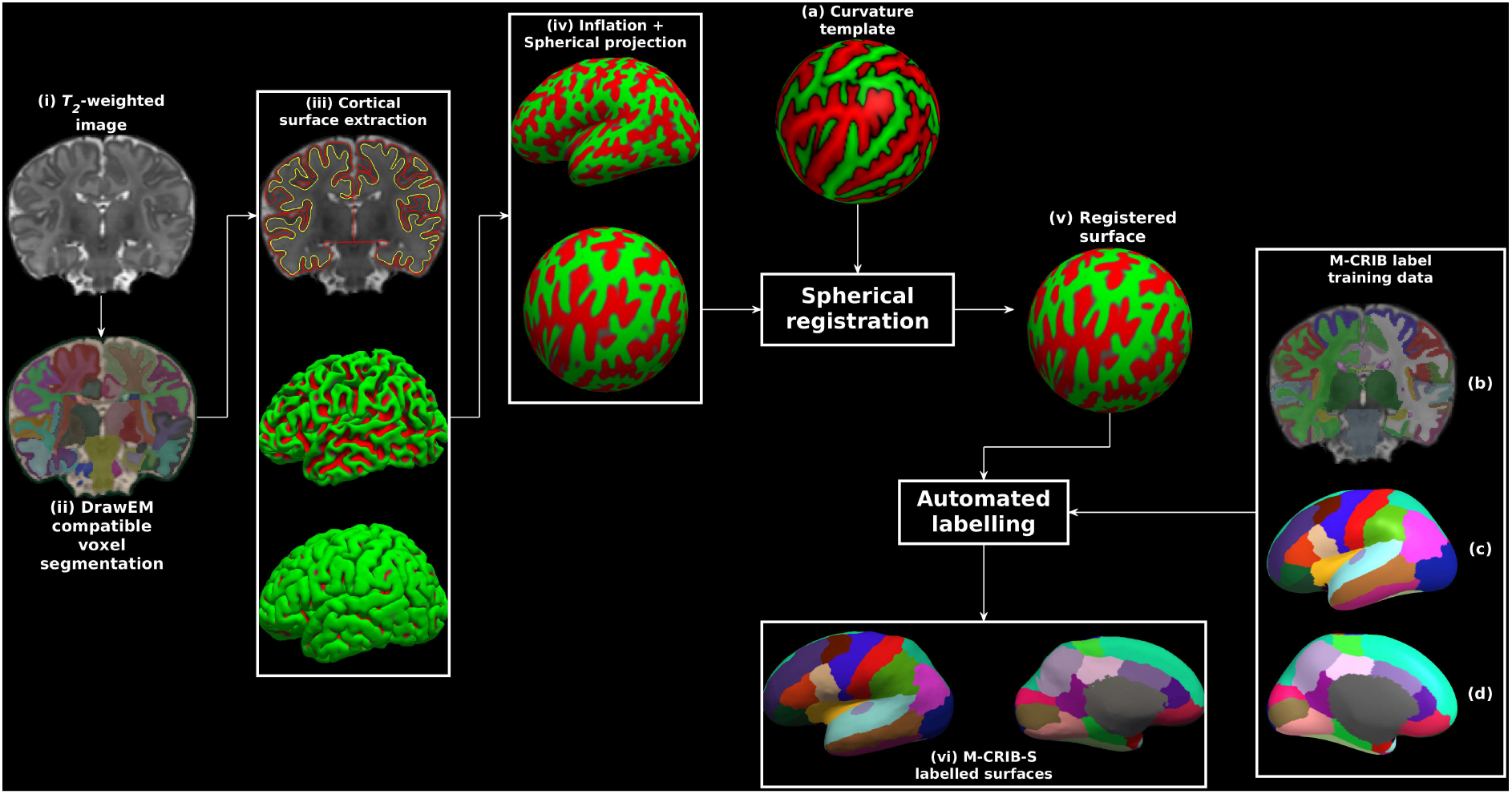
Exemplary surface extraction pipeline output for one labelled subject. Panels show: (i) the original *T*_2_-weighted image; (ii) segmentations according to the *DrawEM* techniques; (iii) *Deformable*-extracted cortical surfaces, where the top panel shows inner (yellow) and outer (red) cortical surfaces overlaid onto the original image, and the middle and bottom panels show lateral aspects of the left hemisphere inner and outer surfaces in 3D; (iv) “inflated” and spherical versions of the white surface; (v) spherical surface registered to the template surface (a); (vi) automatic parcellation using the M-CRIB-S(DKT) scheme shown on the subject inflated surface for lateral (left) and medial (right) aspects. The label training data are depicted in volume format (b), and in 3D on the average inflated surface for lateral (c), and medial (d) aspects. Surface vertices in (iii), (iv) and (v) are coloured according to local mean curvature.

### Image segmentation

Each image in the *unlabelled* dataset (Figure 1(i)) was segmented into cerebral white and grey matter (including lobar sub-divisions), cerebellum and various subcortical grey matter structures automatically using the *DrawEM* software package (Makropoulos et al. 2016; Makropoulos et al. 2014). Briefly, this technique non-linearly registered the non-labelled *T*_2_-weighted images to multiple pre-labelled images. The non-labelled image was then segmented using label fusion. The proposed pipeline utilized the wrapper script neonatal-pipeline-v1.1.sh included in *DrawEM* for execution. Figure 1(ii) shows an example voxel-based *DrawEM* segmentation output.

The *labelled* M-CRIB atlas images were already segmented appropriately for *DrawEM* compatibility. Each M-CRIB segmented image comprised manually traced cerebral white and grey matter, cerebellum, basal ganglia and thalamus, cortical, ventricular and other subcortical labels. Tracing protocols for the cortical (Alexander et al. 2019b; Alexander et al. 2017) and subcortical (Loh et al. 2016) segmentations have been previously described. Figure 1(d) shows an example M-CRIB-S segmented image.

### Surface extraction

*DrawEM* compatible segmentations containing hemispheric white matter and grey matter, cerebellar, ventricular, brainstem and subcortical grey matter labels were used as input for the *Deformable* module (Makropoulos et al. 2018; Schuh et al. 2017) of MIRTK (https://github.com/BioMedIA/MIRTK). *Deformable* used to extract the inner and outer boundaries of each hemisphere of the cerebral cortices for both *labelled* and *unlabelled* datasets. Figure 1(iii)(a) shows inner and outer surfaces overlaid onto the original T2-weighted image, and Figures 1(iii)(b) and 1(iii)(c) show lateral aspects of the inner and outer surfaces in 3D, respectively.

### Surface inflation and spherical mapping

The proposed pipeline used the FreeSurfer tools *mris_inflate* and *mris_sphere* (B. Fischl et al. 1999) to construct inflated and spherical versions of the white matter surfaces, respectively. Figure 1(iv) shows exemplary inflated and spherical surface outputs. Default *FreeSurfer* 6.0.0 options were used for both tools with the following exception: the negative triangle removal option “-remove_negative 1” was added to *mris_sphere*. The inflated surfaces exhibited the same gross shape features as those seen when *FreeSurfer* is executed on adult brain images. Specifically, an overall elliptical appearance, a dimple in the vicinity of the insula, and the smooth protrusion of the temporal and occipital poles.

### Curvature template generation

Surface templates, comprised of all *labelled* and *unlabelled* subjects, were constructed using the curvature-based spherical mapping, alignment and averaging method as previously described (B. Fischl 2012; B. Fischl et al. 1999). Briefly, spherical registration involves linear (rotation) and non-linear displacement of vertices in spherical space. The registration algorithm aims to optimise agreement of white and inflated sulcal depth maps of a subject’s surfaces to a template. The use of local curvatures and sulcal depth to drive registration means that corresponding sulci and gyri are aligned. An iterative procedure of aligning spherical surfaces from both the *labelled* and *unlabelled* datasets to the current template, followed by creation of a new template, was performed. The final template curvature and sulcal depth maps were created by averaging all aligned maps (see Figure 1(a)).

The spherical mapping of each white matter surface onto a common spherical space meant that any given point in template space could be mapped to a point on each subject’s white matter surface, and those points were in correspondence across subjects. This enabled average white, pial and inflated surfaces to be constructed using the *FreeSurfer* tool *mris_make_average_surface*, by resampling surfaces onto the 6^th^ order common icosahedron. The 6^th^ order icosahedron was chosen due to having minimal density while still upsampling the original surfaces.

### Surface labelling

For the 10 cases in the labelled dataset, the volumetric M-CRIB and M-CRIB 2.0 labels were projected to the corresponding white matter surface vertices using nearest labelled neighbour projection. Label data were individually checked for anatomical accuracy of label placement by one author (B.A.). For both atlases, label placement was considered highly accurate. In a few instances, very minor mislabelling was identified and manually corrected on the relevant surface and corrected volumetrically in some cases for M-CRIB 2.0 data.

Figure 2 depicts the projection of the M-CRIB and M-CRIB 2.0 labels projected onto the white matter surface generated by *Deformable* for a single *labelled* subject. These surface-space versions of the M-CRIB 2.0 and M-CRIB parcellations are called M-CRIB-S(DKT) and M-CRIB-S(DK), respectively. While similar, the highlighted regions demonstrate some differences including label boundary changes (in, e.g., lateral orbitofrontal and pars orbitalis) and region removal (banks of the superior temporal sulcus). For a comprehensive description of the differences in regions and region boundaries between the M-CRIB and M-CRIB 2.0 parcellations, see (Alexander et al. 2019b; Alexander et al. 2017).

**Figure 2:**
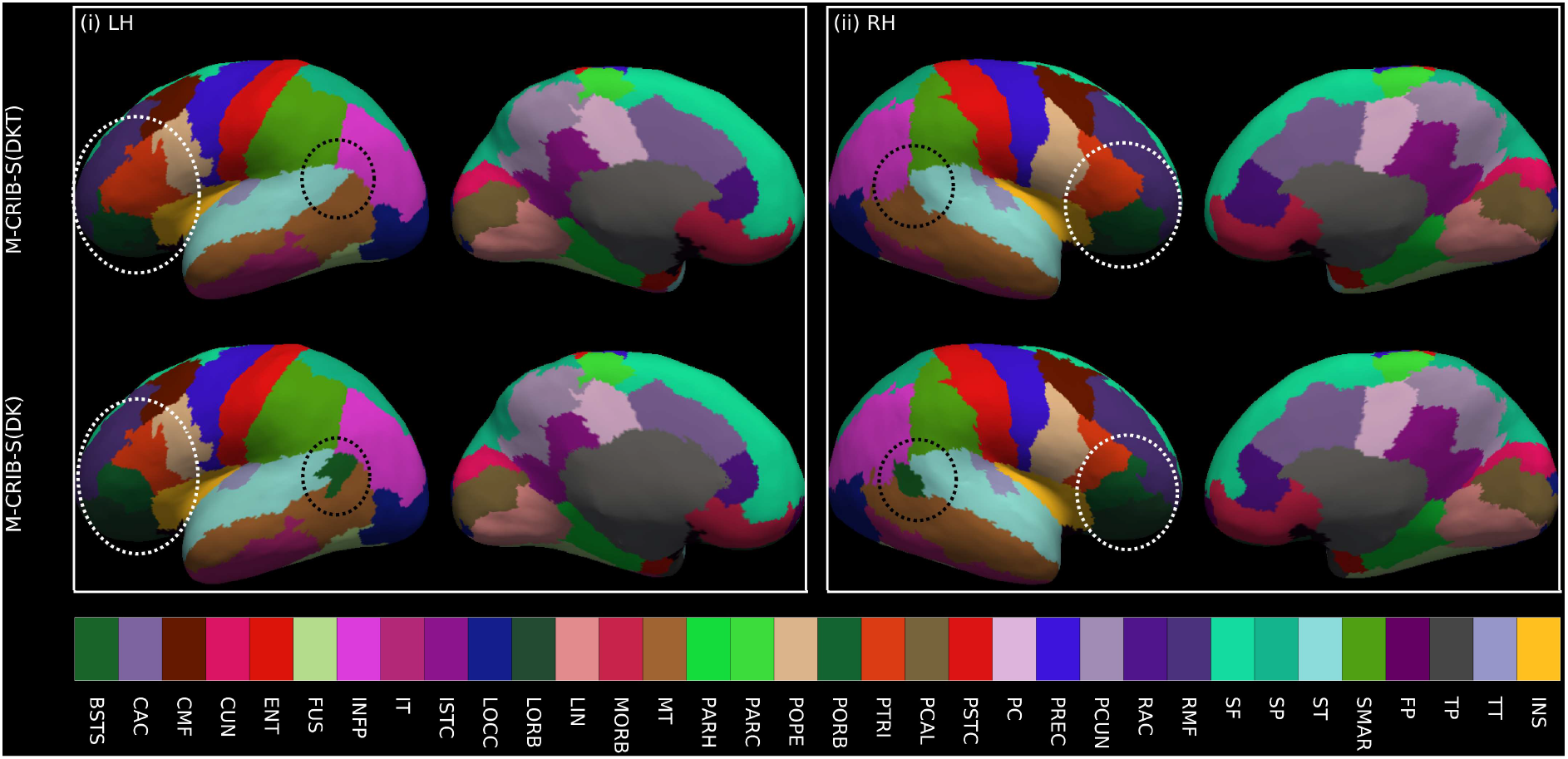
Illustrative surface projections of the manual parcellation for one subject from the *labelled* set using the M-CRIB-S(DKT) and M-CRIB-S(DK) labels for left (LH) and right (RH) hemispheres. The ellipses highlight some differences between M-CRIB-S(DKT) and M-CRIB-S(DK). The white ellipses highlight location disagreements of the lateral orbitofrontal (LORB) and pars orbitalis (PORB) regions between atlases. The black ellipses encompass the banks of the superior temporal sulcus region, which is not present in the DKT.

### Parcellation training set construction

Parcellation training sets were constructed using the *labelled* set for each M-CRIB-S(DKT) and M-CRIB(DK) cortical label, using the method of Fischl et al. (B. Fischl et al. 2004). Briefly, for each template, spatial prior distributions for each cortical label were constructed on the surface using the tool *mris_ca_train*. The M-CRIB-S(DKT) parcellation of the average white surface is shown in Figure 1(c) and 1(d).

### Template surface construction

Using both *labelled* and *unlabelled* datasets, we derived group-averaged white, pial, and inflated surfaces along with curvature and sulcal depth maps in a common spherical space (see Figure 3). For interoperability with the dHCP and UNC atlases (Wu et al. 2018; Makropoulos et al. 2018), we also provide versions of the M-CRIB-S spherical template surfaces registered to the dHCP 42-week and UNC 42-week spherical template surfaces. M-CRIB-S(DKT) and M-CRIB-S(DK) parcellation maps in each *labelled* subject were transferred to the spherical template and used as the training set for the *FreeSurfer* tool *mris_ca_label*. We applied this labelling to the average white matter surface using the M-CRIB-S(DKT) to illustrate our cortical labelling approach (Figures 3(i), (ii) and (iii)). For comparison, the M-CRIB-S(DK) labelling is also shown. These group-average label images may be used for display of statistical analysis results using the M-CRIB-S(DKT) or M-CRIB-S(DK) atlases.

**Figure 3:**
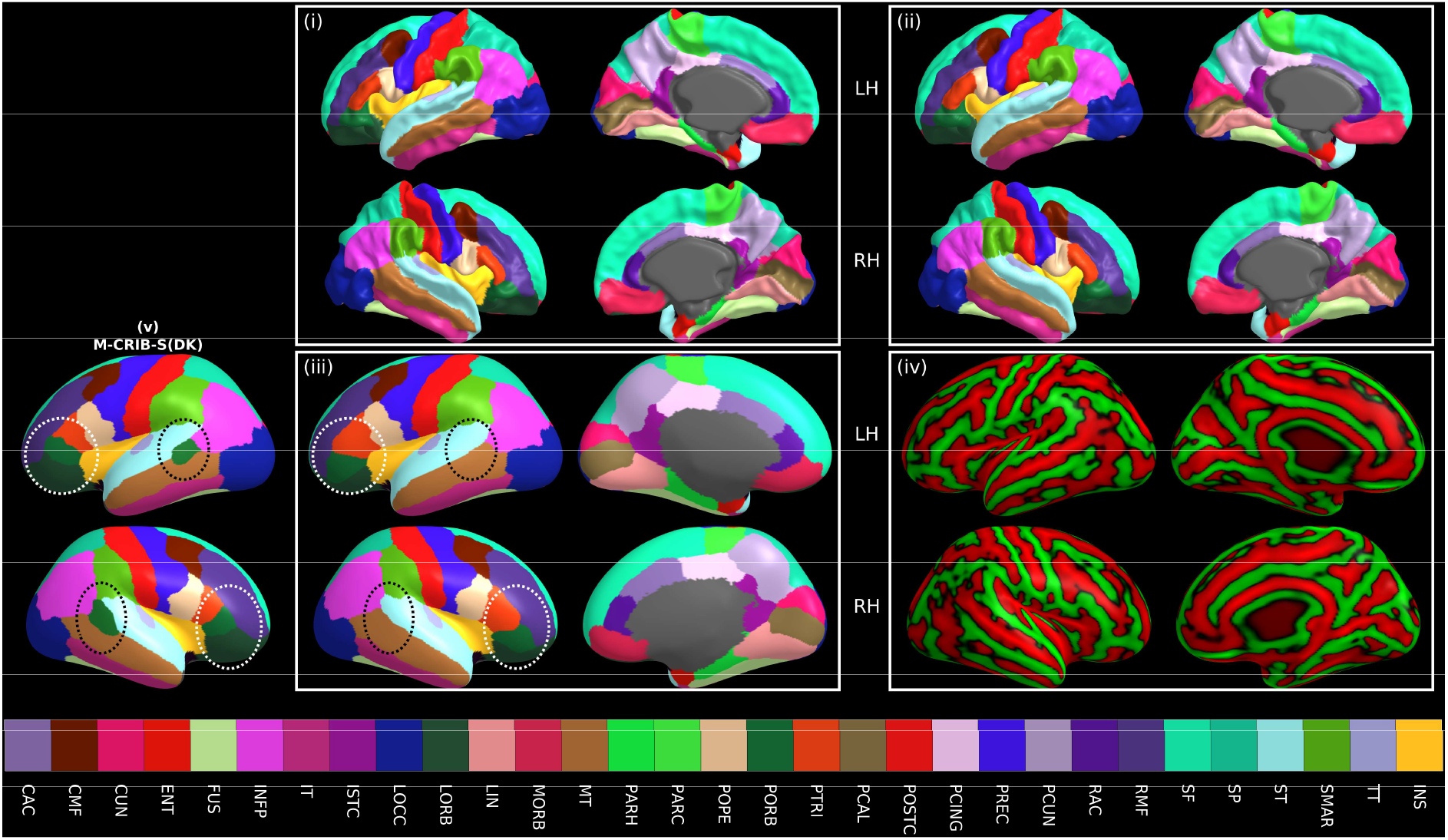
Average white (i), pial (ii), and inflated (iii) surfaces for all subjects with the vertices coloured according to M-CRIB-S(DKT) labels. The average white matter curvature map is shown on the inflated surfaces in (iv). The lateral view of the M-CRIB-S(DK) atlas is shown in (v). The annotations in panel (iii) and (v) highlight selected differences between the atlases. The white ellipses focus on the lateral orbitofrontal and pars orbitalis regions. The black ellipses centre on the bank of the superior temporal sulcus (BSTS), which is absent in the M-CRIB-S(DKT) atlas.

### Parcellation of a novel image

Novel *T*_2_-weighted images can be parcellated using the M-CRIB-S atlas data using the following sequence of processing steps (see Figure 1): 1) Apply *DrawEM* and *Deformable* to extract white and pial surfaces, 2) Perform surface inflation, spherical projection and registration to the M-CRIB-S surface template using FreeSurfer tools, and 3) Use neonatal specific cortical label priors and the automatic labelling tool *mris_ca_label* to parcellate the surfaces. A collection of scripts are provided to execute the pipeline, which can be found along with the M-CRIB-S data on the GitHub page (https://www.github.com/DevelopmentalImagingMCRI/MCRIBS). This pipeline was used to perform cortical parcellation on all *unlabelled* images for validation.

### Parcellation accuracy tests

Parcellation accuracy of the proposed automatic labelling pipeline against manual M-CRIB parcellations was quantified within a Leave-One-Out (LOO) cross-validation framework. For each of the 10 subjects, curvature templates were constructed using the remaining nine *labelled* subjects and all *unlabelled* subjects. Parcellation training data was constructed from the remaining nine *labelled* subjects. The left-out subject was then segmented and parcellated using the pipeline. Per-region label accuracy was assessed using Dice measures, a metric of overlap, and Hausdorff Distances, a metric of boundary error. The Hausdorff distance between the automatic and manual labelling of a region in one subject is the greatest of all shortest distances between two closed contours. Visualisations of these maximal boundary mismatches are provided.

## Results

Results of the Leave-One-Out analysis of labelling accuracy, comparing automated labelling to manually-defined labels in the *labelled* dataset (*n* = 10) are shown in Figures 4, 5 and 6.

**Figure 4:**
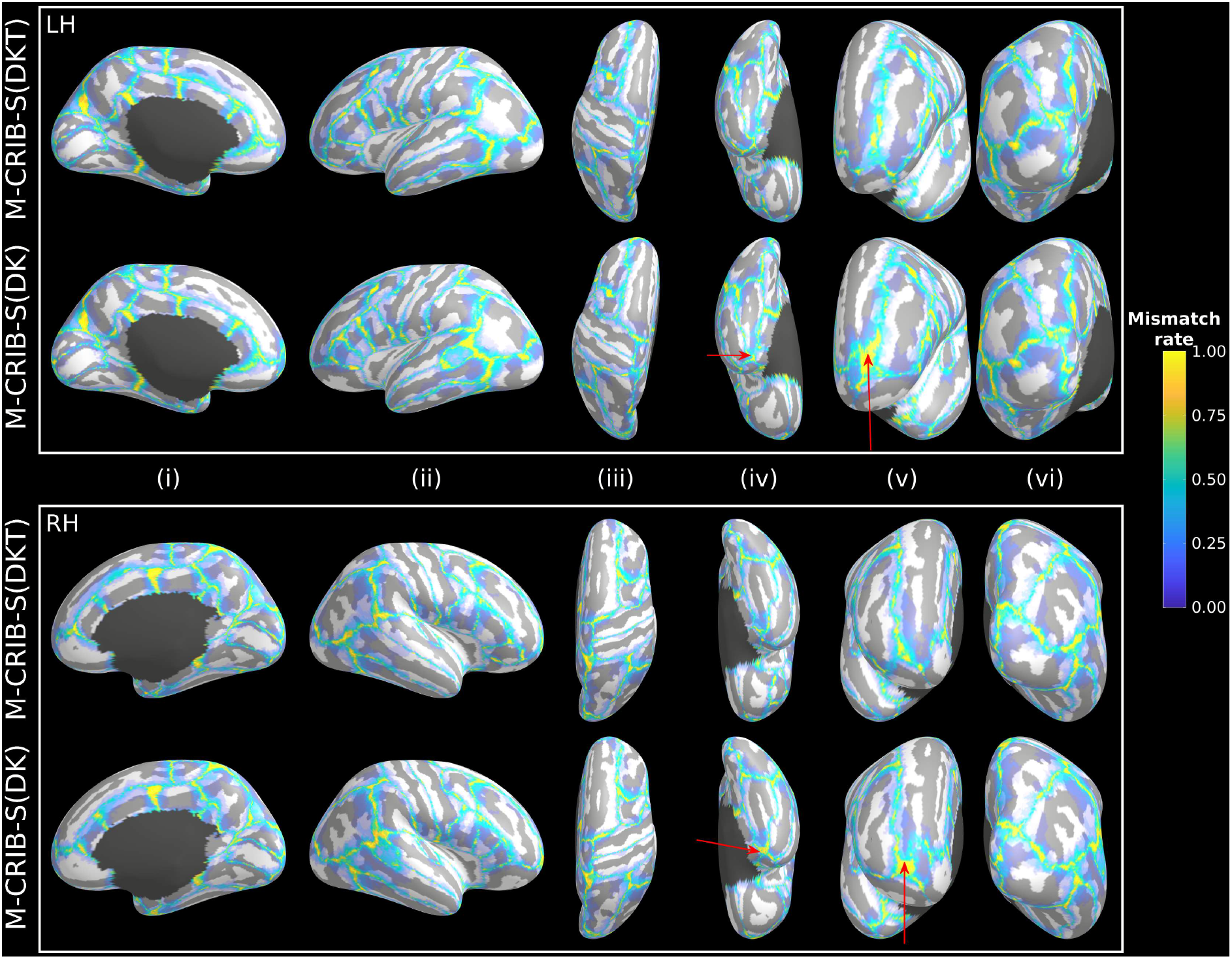
Vertex-wise parcellation mismatch rates for the M-CRIB-S(DKT) and M-CRIB-S(DK) atlases using the Leave-One-Out cross-validation method shown on the template inflated surfaces. Aspects shown are as follows: midline (i), lateral (ii), superior (iii), inferior (iv), frontal (v), and occipital (vi). Warmer colours indicate greater vertex-wise mismatch between automatic and manual labels.

**Figure 5:**
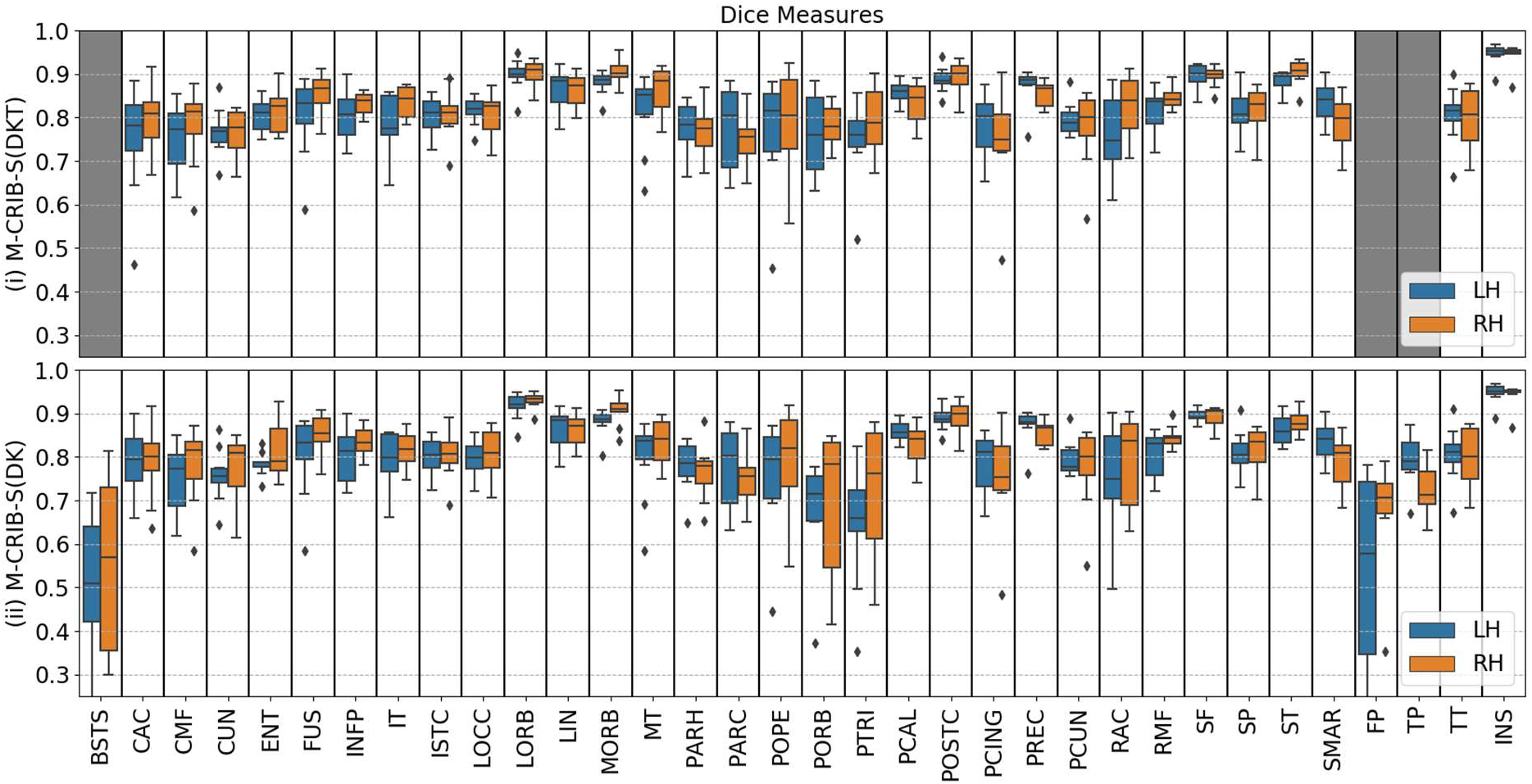
Per-region Dice coefficients for the Leave-One-Out cross-validation test for *labelled* datasets comparing (i) automated with manual M-CRIB-S(DKT) parcellations, and (ii) automated with manual M-CRIB-S(DK) parcellations. The banks of the superior temporal sulcus (BSTS), frontal pole (FP), and temporal pole (TP) regions are greyed out in (i) because they are not present in the DKT parcellation scheme.

**Figure 6:**
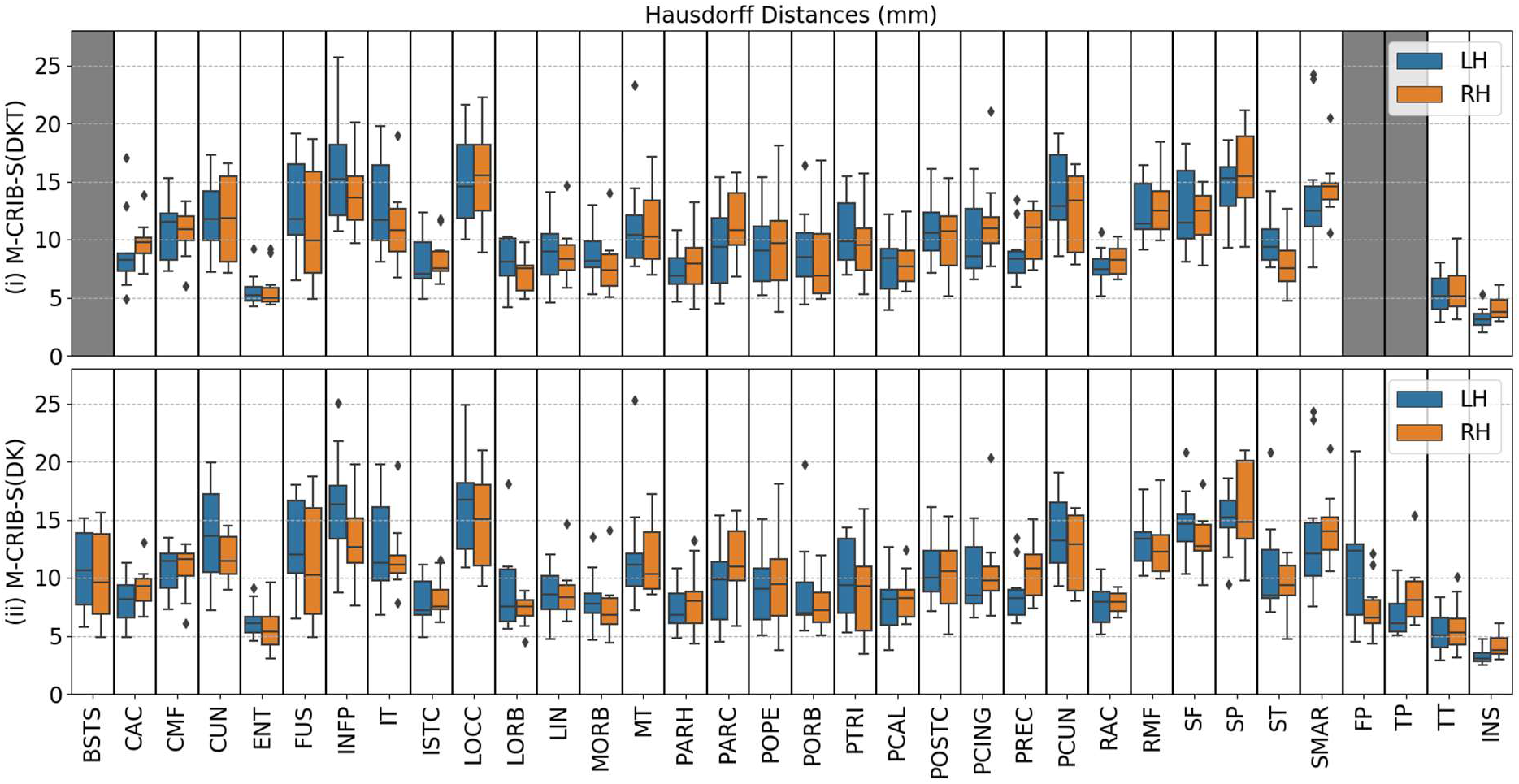
Per-region Hausdorff distances, in mm units, for the Leave-One-Out cross-validation test for *labelled* datasets comparing (i) automated with manual M-CRIB-S(DKT) parcellations and (ii) automated with manual M-CRIB-S(DK) parcellations. The BSTS, FP, and TP regions are greyed out in (i) because they are not present in the DKT parcellation scheme.

Figure 4 shows the vertex-wise parcellation mismatch rates for both atlases. Hemisphere-wide vertex-wise agreement rates were similar across parcellation schemes: the average for M-CRIB-S(DKT) was 0.84 [range 0.78 - 0.87], and the average for M-CRIB-S(DK) was 0.83 [range 0.77 – 0.87]. The rates of high mismatch are confined to region boundaries, indicating that the bulk of the central portions always agreed with ground truth. Exceptionally high rates of mismatch can be seen for the frontal pole and temporal pole labels in the M-CRIB-S(DK).

Figure 5 displays regional Dice measures for M-CRIB-S(DKT) and M-CRIB-S(DK). Dice scores for both atlases were generally high (0.79 - 0.83). For the M-CRIB-S(DKT) parcellation scheme, per-region mean Dice measures were similar across hemispheres (left: mean = 0.82, SD = 0.05; right: mean = 0.83, SD = 0.05). In both hemispheres, the highest Dice scores were observed in the insula (left: 0.95; right: 0.94). The lowest Dice score observed in the left hemisphere was for the pars triangularis (0.75), and the lowest Dice score in the right hemisphere was for the posterior cingulate (0.75). For the M-CRIB-S(DK) parcellation scheme, per-region mean Dice measures were similar to those listed above for the M-CRIB-S(DKT) parcellation, and were again similar between hemispheres (left: mean = 0.79, SD = 0.10; right: mean = 0.80, SD = 0.07). The highest Dices scores in each hemisphere were again seen in the insula (left: 0.95; right: 0.94). The lowest Dice scores were outliers observed in the frontal pole in the left hemisphere (0.50), and in the banks of the superior temporal sulcus region in the right hemisphere (0.55).

Figure 6 shows per-region Hausdorff distances, which measure worst-case boundary discrepancy between automatic and manual labels for M-CRIB-S parcellation schemes. For the M-CRIB-S(DKT) parcellation scheme, per-region mean Hausdorff distances were similar between hemispheres (left: mean: 10.22 mm, SD = 2.99 mm; right: mean: 10.13 mm, SD = 2.77 mm). The smallest Hausdorff distances for each hemisphere were both seen in the insula (left: 3.27 mm, right: 4.11 mm). The largest Hausdorff distance for the left hemisphere was observed in the inferior parietal region (16.18 mm), and the largest in the right hemisphere was for the superior parietal region (15.91 mm).

For the M-CRIB-S(DK) parcellation scheme, Hausdorff distances were similar to those for M-CRIB-S(DKT) and were again similar between hemispheres (left: mean = 10.3 mm, SD = 3.08 mm; right: mean = 9.96 mm, SD = 2.67 mm). The smallest Hausdorff distances observed in each hemisphere were again both in the insula (left: 3.23 mm, right: 4.11 mm). The largest Hausdorff distance seen in the left hemisphere was for the inferior parietal region (16.39 mm), and in the right hemisphere was for the superior parietal region (15.96 mm).

Individual measurements of per-subject and per-label Hausdorff distances ranged from 1.9 - 25.6 mm in M-CRIB-S(DKT) and 2.4 - 25.3 mm in M-CRIB-S(DK). Figure 7 shows some examples of individual worst- and best-case Hausdorff distances between ground truth and estimated labels.

**Figure 7:**
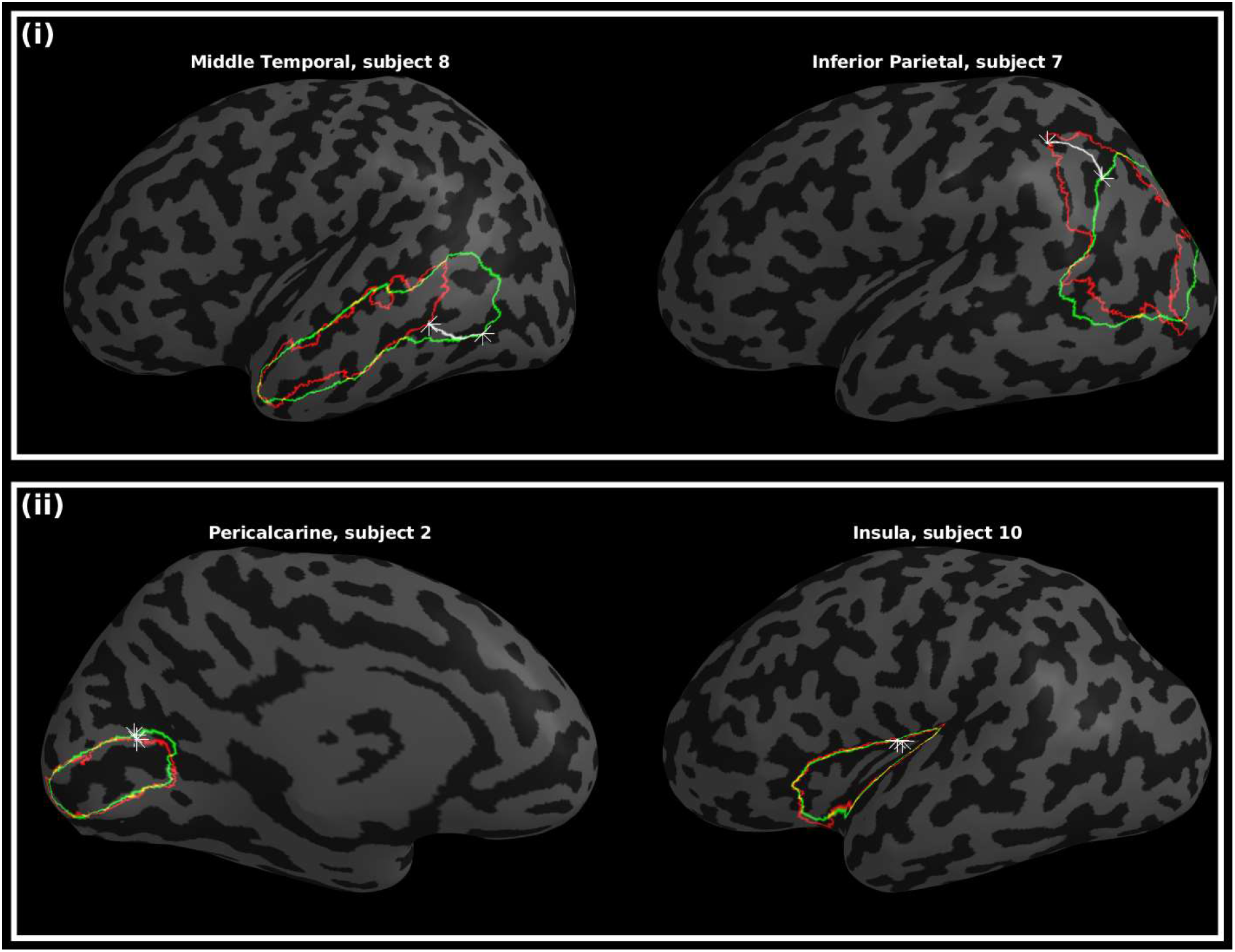
Worst-case (i) and best-case (ii) instances of boundary discrepancies, measured by Hausdorff distances, between estimated (green) and manual (red) label boundaries. The star markers and white paths depict the traversal between nearest neighbours. Other surface vertices are shaded according to curvature, with light and dark grey denoting gyri and sulci, respectively.

## Discussion

The primary contribution of this work is the provision of atlases and tools that facilitate cortical surface extraction and parcellation of the neonatal cortex into 31 or 34 regions. Our pipeline is based on *T*_2_-weighted images of neonates around term equivalent age and uses a common adult-compatible parcellation scheme, with neonatal-specific training data. This work extends our previous M-CRIB and M-CRIB 2.0 volume-based atlases to enable surface-based parcellation of the neonatal cortex.

We have applied this method to a cohort of healthy, term-born infants (mean age at scan = 42.4 weeks). The full age range of subjects suitable for processing under the proposed pipeline will depend on whether tissue intensity contrast is adequate to reliably segment brain structures and extract cortical surfaces, and whether cortical folding complexity is enough to identify all macrostructural morphological features for surface-based template registration and region identification. Tissue segmentation and cortical surface extraction using *DrawEM* and *Deformable* are designed to be compatible with *T*_2_-weighted white/grey matter contrasts visible between 1 - 5 months (Li Wang et al. 2012) and have been demonstrated on data acquired between 34.4 - 43 weeks PMA (Makropoulos et al. 2018). From 5 to 8 months’ PMA the *T*_2_-weighted grey/white matter contrast is transitioning to become like that of an adult by about 9 months’ PMA via an isocontrast phase (6 - 8 months) (Li Wang et al. 2012). As such, tissue segmentation using these protocols would only be expected to work optimally up to approximately 5 months of age, before the *T*_2_-weighted image becomes isointense.

Small-scale cortical folding occurs mostly late in gestation, with many sulci and gyri that define areal boundaries within the M-CRIB-S(DK) and M-CRIB-S(DKT) atlases not being reliably detectable until approximately 40 weeks PMA (Shi et al. 2011). It is possible that a sulcus, for example, may not have formed a sufficiently deep groove to be separated from neighbouring gyri. Thus, careful inspection of parcellation results for subjects scanned at ages below 40 weeks’ PMA is required to ensure that the morphological features that define areal boundaries are present. However, the parcellation component may be appropriate for subjects below 40 weeks and beyond 5 months of age, provided cortical surface data were obtained via other preprocessing methods.

Parcellation accuracy of the automated surface-based labelling methods using the M-CRIB-S atlases was quantified using Dice overlap measures and Hausdorff distances using Leave-One-Out cross-validation. Dice overlap scores were high on average and appeared similar in the left and right hemisphere. The per-vertex mismatch rates (Figure 4) were largely zero for the bulk of the internal portions of most regions. When compared to accuracy measures presented for the adult DKT atlas, Klein and Tourville (2012) presented Leave-One-Out Dice measures of overlap between FreeSurfer parcellations and manual parcellations, which ranged from 0.72 – 0.98. This is largely similar to our Dice overlap results, suggesting that the presented pipeline provides a labelling accuracy consistent with popular adult parcellation tools.

Boundary discrepancies between manual and automated labels were measured using Hausdorff distance. The Hausdorff distance is the maximal distance travelled between any two nearest neighbours of manual and automatic label boundary contours. Most regions had Hausdorff distances between 5 mm - 8 mm for both M-CRIB-S(DKT) and M-CRIB-S(DK) atlases. Figure 7 shows individual instances of worst-case boundary discrepancies in the middle temporal gyrus and inferior parietal labels. The subjects shown, subjects 7 and 8, appeared to exhibit sulcation that varied more than for other subjects in the temporal and parietal cortices and, as a result, the boundaries of these regions were shifted with respect to other training set subjects. An additional confound in manual labelling was that in some instances, label boundaries in the protocols relied on landmarks that were abstract or subject to large individual variability in presence or in morphology. In contrast, the best-case boundary discrepancies (Figure 7(ii)) feature the insula and pericalcarine regions. The high accuracy of the estimated insula boundary is likely due its particularly well-defined boundaries in the original parcellation protocols, relatively easily identifiable in images and consistent across subjects. Other literature has reported cross-validated boundary discrepancy measures between manual and automated segmentations of the adult DK atlas dataset (Desikan et al.). Rather than using Hausdorff distances, discrepancies were calculated as average per-vertex distances between manual and automated label boundaries across subjects. Graphical depictions of these average distances appeared to show a maximum discrepancy of 1 mm. Average boundary mismatch is incompatible with the worst-case discrepancy used in this paper and, thus, cannot be directly compared.

The dHCP set of tools are currently available for cortical surface extraction and lobar parcellation. Makropoulos et al. (2018) recently highlighted the potential value of incorporating M-CRIB parcellation in these tools (Makropoulos et al. 2018), as its compatibility with the DK (Desikan et al. 2006) adult parcellation may facilitate comparisons of cortical measures between the perinatal and adult time points. Here, we provide a publicly available, surfaced based cortical parcellation that can accomplish this objective.

The value of this surface-based atlas and the associated processing scripts is automated parcellation of the neonatal cortex that is straightforward to employ in longitudinal studies. The processing scripts and the M-CRIB-S(DK) and M-CRIB-S(DKT) atlases were constructed to be used with FreeSurfer, to produce compatible output and give a direct correspondence between region-based statistics such as cortical thickness, surface area, and curvature measures at neonatal, childhood and adult timepoints.

## Conclusion

This paper presented the M-CRIB-S(DKT) and M-CBRIB-S(DK) atlases: surface-based versions of the volumetric M-CRIB and M-CRIB 2.0 atlases. It also presented an automated pipeline that involves segmentation of novel *T*_2_-weighted neonatal images, extraction of cortical surfaces, followed by cortical parcellation with the M-CRIB-S(DK) and M-CRIB-S(DKT) atlases, which are neonatal versions of the adult DK and DKT atlases. The curvature template registration targets, average surfaces, labelling training data, and pipeline execution scripts are available. Additionally, for interoperability with the dHCP atlas we have provided a registered version of the spherical template surfaces to be in correspondence to the dHCP template.

## Abbreviations

BSTS: Banks of the superior temporal sulcus
CAC: Caudal anterior cingulate
CMF: Caudal middle frontal
CUN: Cuneus
dHCP: developing Human Connectome Project
DK: Desikan-Killiany
DKT: Desikan-Killiany-Tourville
ENT: Entorhinal
FP: Frontal pole
FUS: Fusiform
INFP: Inferior parietal
INS: Insula
ISTC: Isthmus cingulate
IT: Inferior temporal
L+U: Labelled+Unlabelled
LIN: Lingual
LOCC: Lateral occipital
LOO: Leave One Out
LORB: Lateral orbitofrontal
M-CRIB: Melbourne Children’s Regional Infant Brain
M-CRIB-S: Surface-based Melbourne Children’s Regional Infant Brain atlas
MORB: Medial orbitofrontal
MRI: Magnetic Resonance Imaging
MT: Middle temporal gyrus
PARH: Parahippocampal
PARC: Paracentral lobule
POPE: Pars opercularis
PORB: Pars orbitalis
PCING: Posterior cingulate
PCAL: Pericalcarine
POSTC: Posterior cingulate
PCUN: Precuneus
PREC: Precentral
PTRI: Pars triangularis
RAC: Rostral anterior cingulate
RMF: Rostral middle frontal
SF: Superior frontal
SMAR: Supramarginal gyrus
SP: Superior parietal
ST: Superior temporal gyrus
TP: Temporal pole
TT: Transverse temporal gyrus
UNC: University of North Carolina
PMA: Post-menstrual age
LH: Left hemisphere
RH: Right hemisphere

## Acknowledgements

We gratefully acknowledge support from members of the Victorian Infant Brain Studies (VIBeS) group, Developmental Imaging group, and Melbourne Children’s MRI Centre at the Murdoch Children’s Research Institute, and thank the families who participated in the study. This work was supported in part by the Australian National Health and Medical Research Council (NHMRC) (Project Grant ID 1028822 and 1024516; Centre of Clinical Research Excellence Grant ID 546519; Centre of Research Excellence Grant ID 1060733; Senior Research Fellowship ID 1081288 to P.J.A.; Early Career Fellowship ID 1053787 to J.L.Y.C., ID 1053767 to A.J.S., ID 1012236 to D.K.T.; Career Development Fellowship ID 1108714 to A.J.S., ID 1085754 to D.K.T.), Murdoch Children’s Research Institute Clinical Sciences Theme Grant, the Royal Children’s Hospital, the Department of Paediatrics at the University of Melbourne, the Victorian Government Operational Infrastructure Support Program, and The Royal Children’s Hospital Foundation.

## Notes

#### Summary of Updates

Comparison with UNC atlas removed. Added Hausdorff distance error metric.

